# Synaptic acetylcholine induces sharp wave ripples in the basolateral amygdala through nicotinic receptors

**DOI:** 10.1101/2024.12.01.626291

**Authors:** Joshua X. Bratsch-Prince, Grace C. Jones, James W. Warreni, David D. Mott

## Abstract

While the basolateral amygdala (BLA) is critical in the consolidation of emotional memories, mechanisms underlying memory consolidation in this region are not well understood. In the hippocampus, memory consolidation depends upon network signatures termed sharp wave ripples (SWR). These SWRs largely occur during states of awake rest or slow wave sleep and are inversely correlated with cholinergic tone. While high frequency cholinergic stimulation can inhibit SWRs through muscarinic acetylcholine receptors, it is unclear how nicotinic acetylcholine receptors or different cholinergic firing patterns may influence SWR generation. SWRs are also present in BLA *in vivo*. Interestingly, the BLA receives extremely dense cholinergic inputs, yet the relationship between acetylcholine (ACh) and BLA SWRs is unexplored. Here, using brain slice electrophysiology in male and female mice, we show that brief stimulation of ACh inputs to BLA reliably induces SWRs that resemble those that occur in the BLA *in vivo*. Repeated ACh-SWRs are induced with single pulse stimulation at low, but not higher frequencies. ACh-SWRs are driven by nicotinic receptors which recruit different classes of local interneurons and trigger glutamate release from external inputs. In total, our findings establish a previously undefined mechanism for SWR induction in the brain. They also challenge the previous notion of neuromodulators as purely modulatory agents gating these events but instead reveal these systems can directly instruct SWR induction with temporal precision. Further, these results intriguingly suggest a new role for the nicotinic system in emotional memory consolidation.

**Significance Statement:** Sharp wave ripples represent a key network signature believed to be important in memory consolidation. These network events have been largely studied in the hippocampus and are suppressed by the basal forebrain cholinergic system. Sharp wave ripples also occur in the basolateral amygdala, a region critically involved in emotional memory consolidation. Here, we show a novel mechanism by which low frequency stimulation of synaptic acetylcholine in basolateral amygdala brain slices can reliably induce sharp wave ripples through a nicotinic receptor-mediated mechanism. These findings update the view of neuromodulatory systems in sharp wave ripple induction, moving past a purely modulatory role for these systems and instead establish a role where modulators, like acetylcholine, can directly drive SWRs with temporal precision.

## Introduction

Memory consolidation requires the proper storage of past experiences in a manner that is available for reactivation. In the hippocampus, these processes are closely associated with network events termed sharp wave ripples (SWR), consisting of a large amplitude, slow frequency (1-20 Hz) envelope (sharp wave, SW) with a brief high-frequency oscillation (150-200 Hz ripple) that exhibit strong similarities in both human and rodents (1–3). Hippocampal SWRs predominantly occur during periods of immobility and slow wave sleep (SWS) (4) and replay precise sequences of past neuronal activity (5, 6). SWRs are increased after learning (7), can induce plastic changes in cell assemblies (8, 9), and when disrupted can block memory formation (10), positioning them as a neural substrate for memory consolidation.

Emotional arousal often leads to enhanced memory consolidation and stronger memories. The basolateral amygdala (BLA) plays a driving role in emotional memory consolidation (11), such as associative fear learning, a critical function for animal survival. Plasticity within populations of neurons in the BLA are observed after learning (12, 13) and when reactivated can induce memory recall (14). However, mechanisms underlying BLA ensemble consolidation are not clear. SWR activity has been reported in the BLA *in vivo* (15) and represents a mechanism by which BLA ensembles could undergo memory consolidation. Similar SWR profiles have been observed in *in vitro* preparations in the BLA (16–18). Despite this potential role in BLA memory consolidation, a mechanistic understanding for BLA SWRs is lacking, yet offers valuable insight into how memories of emotionally charged experiences are consolidated.

The exact circuit and synaptic mechanisms by which SWRs arise are still being uncovered. In the hippocampus, *in vivo* studies have outlined a role for synchronized discharge of a population of CA3 pyramidal neurons (PYRs) which depolarize the apical dendrites of CA1 PYRs (producing a SW) and produce ripple firing among populations of these cells (1, 19). Different classes of local interneurons (INs) fire spikes at distinct times in relation to the ripple (20) and are likely recruited by local PYR-IN interactions. *In vitro* preparations can also spontaneously exhibit SWRs that share characteristics with those *in vivo* and offer insight into the synaptic mechanisms underlying SWR generation (21). These studies have established that SWRs arise from precise interactions of excitation and inhibition. Accordingly, neuromodulatory systems, which regulate excitation and inhibition have been shown to impact SWR generation. Hippocampal SWRs largely occur during periods of immobility and SWS, which inversely correlates with cholinergic tone (22, 23). Consistent with this, cholinergic stimulation at high frequencies, a stimulation pattern reflecting cholinergic neuron activity during active exploration or REM sleep, or application of cholinergic agonists have been shown to inhibit SWRs through a muscarinic receptor-mediated mechanism (24–28). This positions basal forebrain neurons through acetylcholine (ACh) signaling to gate network states permitting SWR generation.

Interestingly, the BLA receives some of the densest cholinergic projections from the basal forebrain (29), but the relationship between ACh and BLA SWRs is not clear. Further, it is unexplored how low frequency firing of cholinergic neurons, as is seen during immobility or SWS, impacts SWRs. ACh can act through both through ionotropic nicotinic receptors and metabotropic muscarinic receptors on different timescales to differentially impact circuit function (30–32). Compared to muscarinic receptors, nicotinic receptors act on a much faster, within milliseconds, timescale that is more aligned with SWR kinetics. Nicotinic receptors can also be engaged by lower concentrations of ACh than muscarinic receptors, positioning the nicotinic system to exhibit larger influence during periods of lower cholinergic tone. In total, our understanding of how the basal forebrain cholinergic system impacts SWRs is incomplete. In this study, we show that low frequency stimulation of cholinergic fibers in the BLA can induce local field potential (LFP) SWRs through a nicotinic receptor mechanism that has not yet been implicated in SWR generation. These findings expand our knowledge of mechanisms generating SWRs and suggest intriguing possibilities through which the basal forebrain cholinergic system can regulate emotional memories.

## Results

### Single pulse optogenetic stimulation of cholinergic fibers induces sharp-wave ripple activity in the BLA

To explore the effects of synaptically released ACh on BLA network activity, we first performed LFP recordings in the BLA in coronal brain slice preparations of ChAT-ChR2 mice (Fig. 1A). Measurable baseline LFP activity in BLA slices was low, consistent with the activity of the amygdala *in vivo,* but did show periodic (<0.1 Hz) short bursts of activity (Fig. 1B). Analysis of these events revealed distinct frequency components: a large amplitude downward slow wave event that was accompanied at its peak by a short duration, high frequency event, consistent with SWRs in slice preparations (18). Interestingly, single pulse light stimulation (490 nm, 1-2 ms duration) of cholinergic fibers produced a quick onset network event (Figure 1B, right) that showed similar characteristics: a slow wave accompanied by a high frequency activity at its peak. Spectrogram analysis of this light-induced event revealed the high-frequency component peaked around 160-200 Hz; a frequency consistent with that of ripples. While almost all slices exhibited the slow frequency event in response to light, the high frequency event occurred in just over half of these slices tested (55%; 12/22 slices (n), 15 animals (N)). Overall, this event is consistent with amygdalar SWRs observed both *in vivo* (15) and *in vitro* (18). We termed these events as ACh-SWRs.

**Figure 1.**
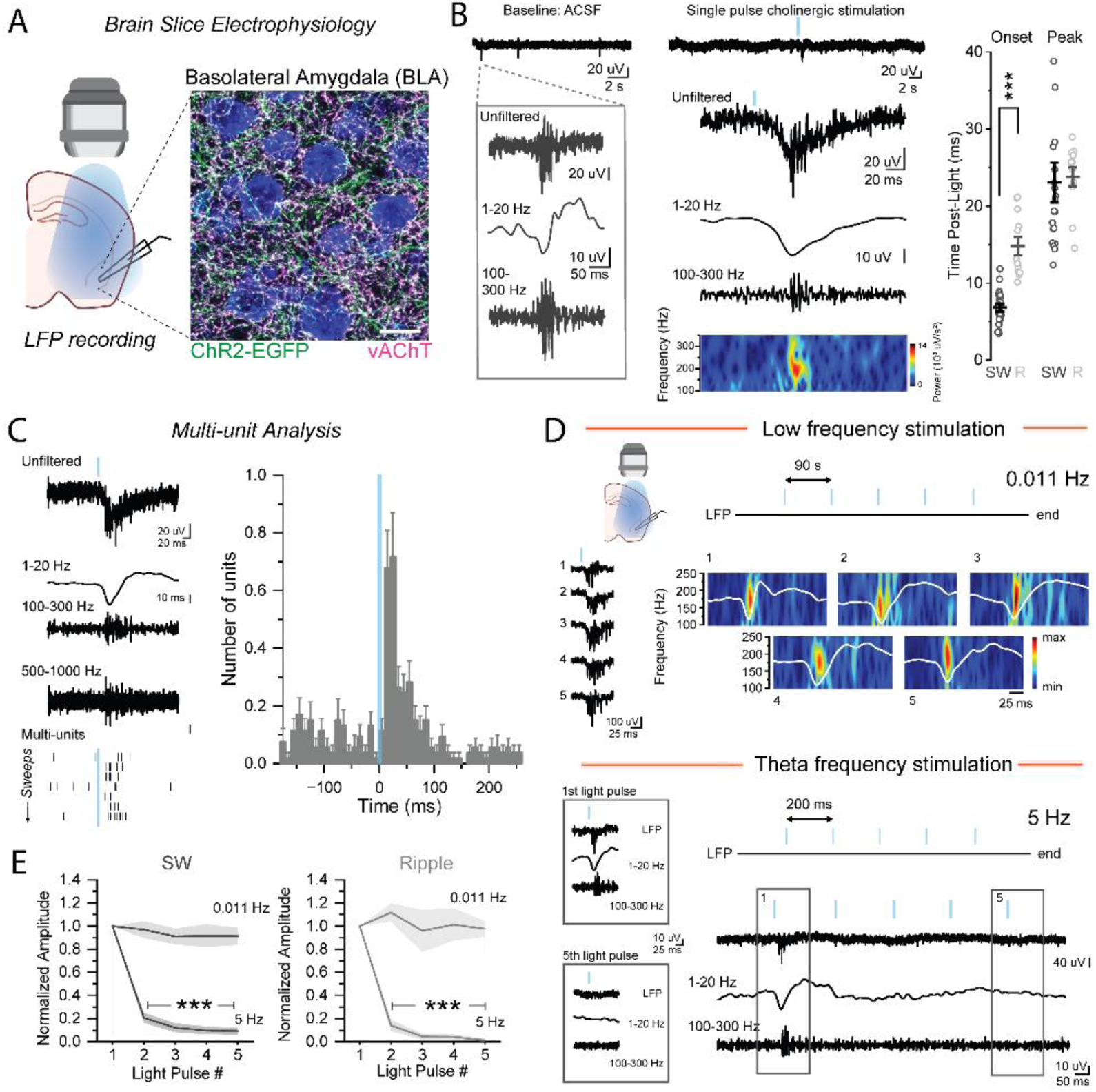
Single pulse stimulation of cholinergic fibers induces sharp wave ripples in BLA slices. **A**. A confocal image of a slice of the BLA of a ChAT-ChR2 mice showing dense vAChT+ (magenta) terminals that co-express ChR2 (green). Nissl stain shown in blue. **B**. In baseline, LFP recordings show periodic events displaying an overlapping slow wave (filtered at 1-20 Hz) and high frequency ripple (filtered at 100-300 Hz) (left). Single pulse stimulation of ACh fibers induces an event with similar frequency characteristics, termed ACh-SWR (right). Analysis of the timing of these events shows that the sharp wave (SW) precedes that of the ripple (R) while both peak at similar times. **C**. Isolation of multiunit activity from these recordings show that unit activity is increased during this ACh-SWR. **D.** During low frequency stimulation (0.011 Hz, top), ACh-SWRs are reliably induced on each light pulse (traces, left). Spectrograms of each light pulse show a similar frequency and timing of the ripple in relation to the filtered slow wave (white trace). During theta frequency stimulation (5 Hz), ACh-SWRs are only induced on the first light pulse. Traces on the bottom left (gray square) highlight the differences in the first and fifth pulse. **E.** Comparing the amplitude of the SW (trace filtered at 1-20 Hz) and the R (filtered at 100-300 Hz) of each light pulse at 0.011 and 5 Hz shows that these events are reliably produced with each stimulation only at the lower frequency, with a significant decrease in both of these features on subsequent pulses at 5 Hz.

Examining the time course of ACh-SWRs also revealed strong similarities to both hippocampal and amygdala SWRs *in vivo*. The slow, sharp wave (SW) component had a remarkably fast onset following light stimulation that preceded the ripple (SW onset from start of light pulse: 6.8 ± 0.4 ms, n= 22, N = 15; ripple onset: 14.8 ± 1.2 ms, n = 12, N = 12; paired t-test, p = 1.73 x 10^-4^) while both peaked at similar times (SW = 23.3 ± 2.5 ms; ripple = 24.0 ± 1.2 ms; paired t-test, p = 0.84). The entire ACh-SWR duration ranged between 60–125 ms (average: 98.9 ± 6.6 ms) and was often accompanied by a slow rebound upward event upon completion, as is observed in the hippocampus. There was no measurable difference in induced SWRs in male and female mice including SW amplitude (male (M) = 0.023 ± 0.002 uV, n =14, N = 10; female (F) = 0.022 ± 0.002 uV, n = 8, N = 5; two sample t-test, p = 0.91), SW duration (M = 98.74 ± 9.49 ms; F = 99.28 ± 7.81 ms; two sample t-test, p = 0.97), ripple incidence (M = 57%, 8/14 slices; F = 50%, 4/8 slices) or ripple duration (M = 30.09 ± 2.65 ms, n = 8, N = 6; F = 23.88± 5.45 ms, n = 4, N = 2; two sample t-test, p = 0.27).

In both the hippocampus and amygdala, SWRs are associated with increased spiking in a small portion of PYRs (15, 19, 33). To explore if the same occurs with ACh-SWRs, we measured multi-unit spiking activity during ACh-SWRs (Figure 1C). Indeed, low multi-unit activity in baseline was robustly increased during the ACh-SWR directly following light stimulation (average of 53 SWR events, N = 14). Peak increases in unit frequency occurred near the peak of the ACh-SWR and were short in duration, returning to baseline levels within 100 ms after light stimulation. This indicates that BLA ACh-SWRs are also associated with an increase in spiking activity, as is observed *in vivo*.

### Cholinergic induction of SWRs in the BLA only occurs with low frequency cholinergic stimulation

In the hippocampus, SWR occur periodically, yet irregularly, during immobility and slow-wave sleep at a rate of 2-30/minute (4, 34), rates that extend to SWRs in the BLA (15). We were next interested in the ability of repeated cholinergic stimulation to induce SWRs in the BLA. When single pulse light stimulation was given every 60-90 seconds, there was a surprisingly remarkable consistency in the ACh-SWR (Figure 1D, stimulation frequency 0.11Hz). The induced ripple in a given preparation showed a similar repeated peak frequency and occurred at a similar time in relation to the large SW that could persist for over 30 minutes of this low frequency stimulation. Interestingly, however, repeated light stimulation at theta frequency (∼5Hz), a frequency at which basal forebrain cholinergic neurons fire *in vivo* during an awake and alert state (22), did not result in repeated ACh-SWRs on each light pulse (Figure 1E). While the first light pulse induced a ACh-SWR, subsequent light pulses produced increasingly small SWs and no obvious detected high frequency ripple events (Fig. 1E: 5^th^ light pulse: normalized SW amplitude = 9.0 ± 2.9% of 1^st^ light pulse; one way repeated measures ANOVA, p = 7.78 x 10^-11^, F(4,11) = 263.00, n = 15 slices, N = 9; normalized ripple amplitude = 1.4 ± 1.0% of 1^st^ light pulse, one way repeated measures ANOVA, p = 4.05 x 10^-7^, F(4,4) = 2717.51; n = 8 slices, N = 6). We have previously shown in these preparations that ACh is reliably released at this frequency (35), ruling out a depletion in the cholinergic fibers as the causing factor. These results indicate that periodic low frequency synaptic ACh stimulation can reliably evoke ACh-SWRs in the BLA. However, theta frequency stimulation does not produce continued ACh-SWRs with each light pulse.

### Nicotinic acetylcholine receptors drive SWRs in the BLA that are dependent on both glutamatergic and GABAergic mechanisms

We were next interested in which type of ACh receptors were involved in ACh-SWR induction and how glutamatergic and GABAergic systems in the BLA were recruited in these events. To systemically explore this, we broke down the ACh-SWR into its different components: the SW, ripple, and unit activity, and probed the receptors involved in each. Isolation of the SW after light stimulation revealed no effect on SW amplitude by the general muscarinic receptor antagonist atropine (atro., 5 uM; 0.922 ± .062 of control; n = 8, N = 6; paired t-test, p = 0.24). Instead, a complete reduction was observed by the general nicotinic receptor antagonist mecamylamine (mec., 10 uM; 0.090 ± 0.034 of control; n = 11, N = 7; paired t-test, p = 1.31 x 10^-4^), indicating that the SW was a nicotinic receptor-mediated event (Figure 2A). To next determine if there was any contribution of glutamatergic or GABAergic systems to the SW generation, application of the AMPA receptor antagonist CNQX (20 uM) or the GABA_A_ receptor antagonist picrotoxin (40 uM) were applied in a separate set of experiments (Figure 2B). Blocking AMPA receptors resulted in a significant decrease in SW amplitude (CNQX: 0.576 ± 0.079 of control; n = 7, N = 4; paired t-test, p = 0.024) whereas blocking GABA_A_ receptors resulted in a smaller, yet still significant, reduction of SW amplitude (picro: 0.751 ± 0.083 of control; n = 6, N = 4; paired t-test, p = 0.028). Taken together, these results indicate that the SW component of the ACh-SWR were evoked by nicotinic receptors that recruited AMPA and GABA_A_ signaling.

**Figure 2.**
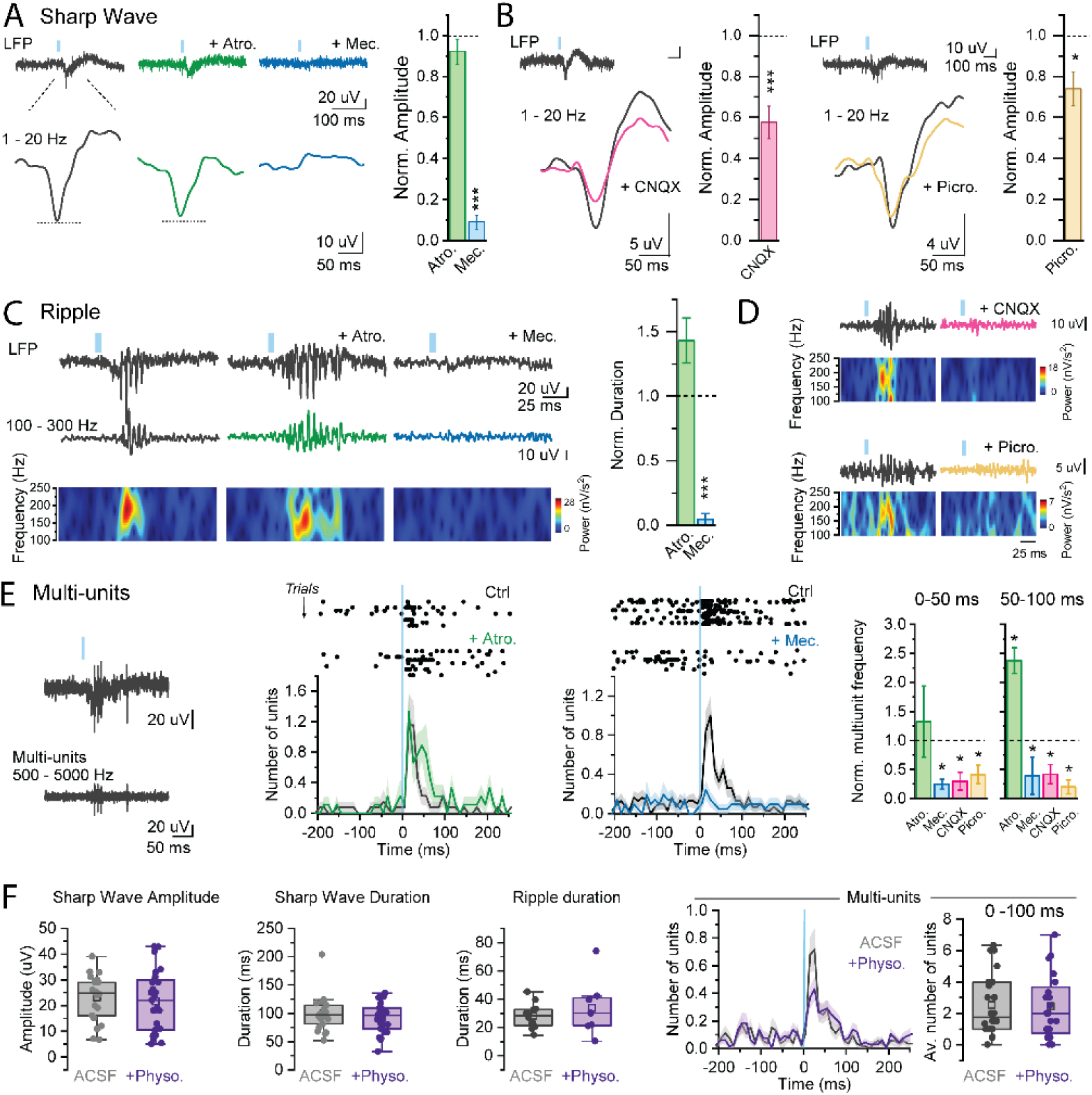
Acetylcholine acts through nicotinic receptors to drive BLA sharp wave ripples and recruits local glutamatergic and GABAergic systems. **A.** Isolation of the sharp wave of the LFP (filtered at 1-20 Hz) shows that its amplitude is not affected by atropine (5 µM), but the sharp wave is blocked by mecamylamine (10 µM) **B.** Application of CNQX (20 µM) and picrotoxin (40 µM) reduces sharp wave amplitude but does not completely block it. **C.** Isolation of ripple activity (filtered at 100-300 Hz) shows that atropine does not block but extends ripple duration. In contrast, ripple activity is blocked by mecamylamine, as quantified by ripple duration. **D**. Example of ripple filtered traces and spectrograms show that application of CNQX (top) or picrotoxin (bottom) completely blocks ripple activity in the LFP. **E.** Multi-unit quantification shows that atropine application increases unit firing in the 50-100 ms time window following cholinergic stimulation while increased multi-unit activity during the SWR is completely blocked by mecamylamine (top: raster plots of example slice; bottom: average frequency across all slices). Application of CNQX or picrotoxin also strongly reduces the number of units following light stimulation (right). **F.** Application of physostigmine (1 uM) has no effect on sharp wave amplitude, sharp wave duration, or ripple duration. Additionally, physostigmine does not change the immediate (0-100 ms post-light) increase in multiunit frequency induced by ACh stimulation.

We next explored which receptors were involved in the high frequency ripple generation (Figure 2C). Consistent with the SW, application of atropine did not block the induction of the ripple in any of the slices (n = 5) but instead resulted in a trend of increased ripple duration (atro. = 1.43 ± 0.17 of control; pair sample t test, p = 0.13). Application of mecamylamine completely blocked ripple activity (mec. = 0.04 ± .04 of control; paired t test, p = 0.002) indicating that ripple generation, in addition to the SW, were induced by nicotinic receptor signaling. Further, both CNQX and picrotoxin application completely blocked ripple generation (Figure 2D), indicating a critical role of glutamatergic and GABAergic systems in ripple generation. Finally, these pharmacological results extended to BLA multiunit activity. Atropine did not block the increase in light-induced multiunit activity but instead prolonged the duration of this increased activity, specifically in the 50-100 ms window following light stimulation (Figure 2E). Meanwhile, mecamylamine blocked the increase in unit activity by ACh. Consistent with the close relation of multi-unit activity to the ripple, blockade of both AMPA and GABA_A_ receptors also significantly reduced the increase in multi-unit activity (Figure 2E). Interestingly, application of low levels of the acetylcholinesterase inhibitor physostigmine (1 uM), prolonging the effects of ACh in the brain slice, did not affect any SWR properties (Figure 2F) or increase the incidence of ripple generation. In total, these results show that cholinergic stimulation induces ACh-SWRs in the BLA strictly through nicotinic receptors and the recruitment of local glutamate and GABA systems. Further, they indicate that muscarinic receptor activation suppresses SWR generation, constraining the event to a discrete temporal window following cholinergic stimulation.

### Whole cell correlates of ACh-SWRs reveal coordinated nicotinic-mediated sEPSC and sIPSC recruitment

To get a mechanistic understanding of the ACh-SWR at the synaptic level, intracellular whole-cell patch clamp recordings were performed. During simultaneous LFP and voltage clamp whole cell recordings of BLA PYRs, large compound events were observed in PYR recordings that occurred during the LFP SWR (Figure 3A). Holding the PYR at -70 mV (at Cl-reversal for GABA_A_ receptors) revealed a barrage of inward currents, while holding at 0 mV (at AMPA reversal) produced a large barrage of outward currents, suggesting involvement of both AMPA and GABA_A_ events. Examining the time course for the EPSC and IPSC onset after ACh stimulation revealed that EPSC onset preceded that of the IPSC (Fig 3B; EPSC: 4.23 ± 0.09 ms after light pulse; 27 cells (n), N = 10; IPSC: 8.07 ± 0.89 ms; n = 22, N = 15; two sample t-test, p = 1.97 x 10^-5^) while both the EPSC and IPSCs peaked at similar times (EPSC: 17.15 ± 1.38 ms; IPSC: 21.39 ± 1.93 ms; two sample t-test, p = 0.07), a time course that is similar to the ACh-SWR of the LFP. In recording of neighboring PYRs, it could be shown that EPSC activity preceded that of IPSC within the same ACh-SWR (Figure 3B). A closer look at both the EPSC and IPSC events in BLA PYRs revealed they consisted of a summation of many events that could be identified by peak detection. On average, EPSCs consisted of around 4 events (3.9 ± 0.3) with an inter-event interval of 9.61 ± 0.85 ms and IPSCs consisted of almost 4.5 events (4.4 ± 0.3) at an average inter-event interval of 6.3 ± 0.4 ms. Again, the rate of these EPSCs and IPSCs align with the frequency range of the LFP ripple. Interestingly, in repeated trials within the same cell (stim frequency of 0.011Hz), there was a great consistency in the timing of the peaks of both EPSCs and IPSCs (Figure 3C, example cell event probability in 2ms time bins) induced by ACh stimulation, consistent with what was observed at the level of the LFP SWR. This suggests that ACh stimulation is reliably producing a similar network event on each trial with impressive temporal repeatability.

**Figure 3.**
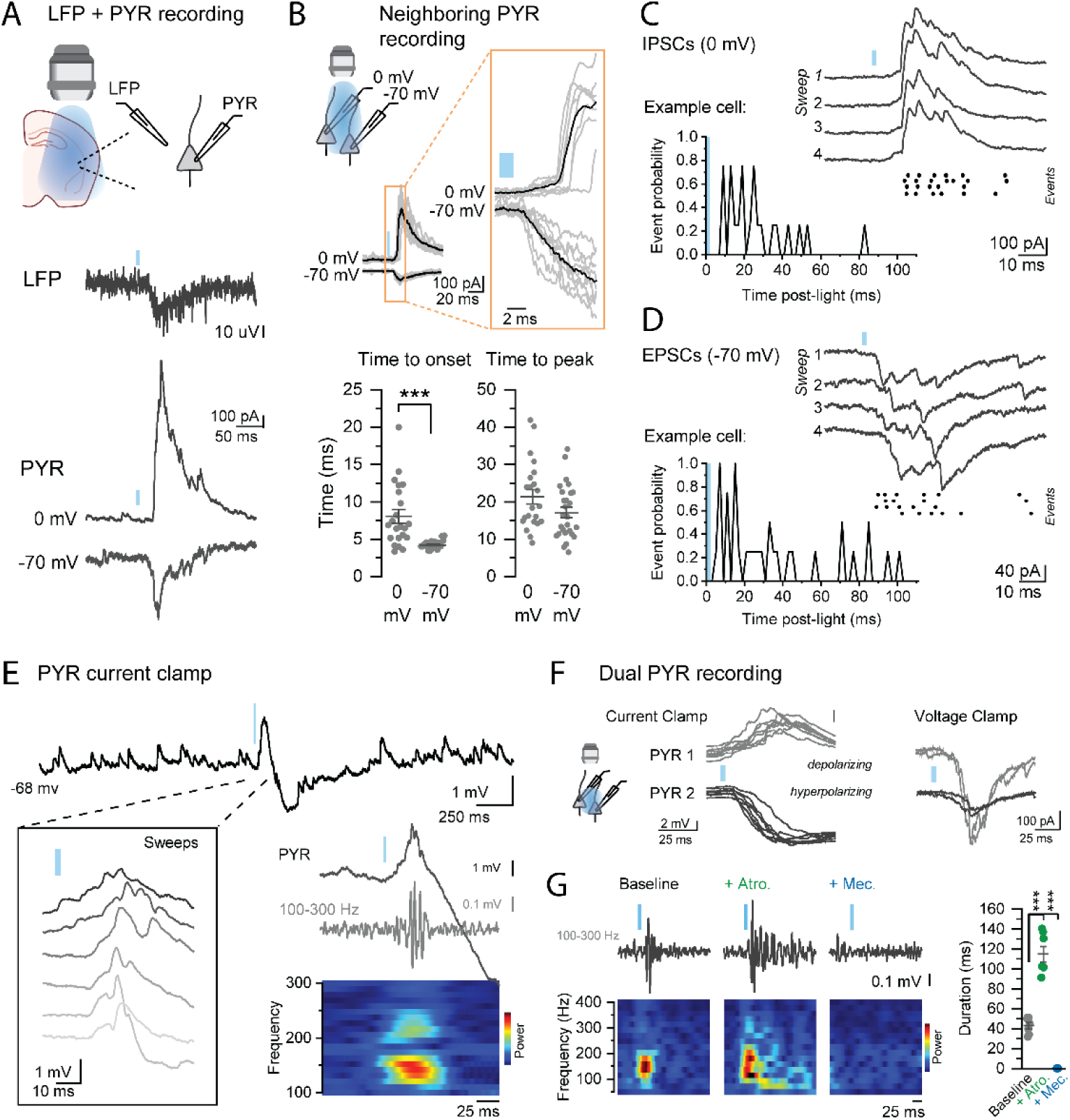
Acetylcholine stimulation induces a barrage of excitatory and inhibitory currents in BLA PYRs that are driven by nicotinic receptors. **A**. Example traces of simultaneous LFP and whole cell patch clamp recording (PYR) showing a response at the single cell level during ACh-SWR of the LFP that differs when the cell is held at 0 or -70 mV. **B**. Example recording of dual PYRs where one cell is held at 0 mV and one at -70 mV shows that the inward current event at -70 precedes the outward current event at 0 mV in the same slice preparation. Comparing onset and peak responses of these events across all cells shows that while they peak at similar times, the event at -70 mV precedes that of the event at 0 mV. **C,D.** Isolation of the IPSC and EPSC events at 0 mV and -70 mV, respectively, show that they consist of a summation of smaller events that can show remarkable consistency across ACh stimulations (waveforms showing example cells). Histograms show the timing distribution of these events of example cells across trials occur with repetitive temporal precision. **E**. Current clamp recordings of BLA PYRs show a ripple frequency in the membrane potential (filtered trace and spectrogram, right) that is also repeated across ACh stimulations (multiple sweeps overlayed, left). **F.** Example current clamp traces from dual PYR recordings at a resting membrane potential, where one PYR shows a depolarizing response, and one shows a hyperpolarizing response during the ACh-SWR. Voltage clamp recordings of these same PYRs show a large difference in the amplitude of the fast EPSCs evoked by ACh that aligns with the membrane potential changes. **G.** Example traces filtered at 100-300Hz from a whole cell PYR current clamp recording show an increased duration of ripple frequency activity with application of Atro. and complete blockade by Mec. (quantification of all cells on right).

To explore how these synaptic events influence PYR dynamics, we also performed current clamp recordings of these cells at rest during ACh stimulation (Figure 3E). At a resting membrane potential, ACh stimulation evoked an immediate ripple-frequency membrane oscillation that was consistent across multiple trials in the majority of PYR recordings. In all cells, this was followed by a slow hyperpolarization, as has been previously shown in BLA PYRs (36, 37). As we had noticed a large range in the amplitude of nicotinic mediated EPSCs and IPSCs (EPSC peak amplitude range 10-500 pA, IPSC peak amplitudes from 100-2000 pA), we hypothesized that this range could lead to differences in the overall response across PYRs. Indeed, while some cells were depolarized during this ripple event, others showed a hyperpolarizing response. During dual PYR recordings, it was possible to observe both types of responses in neighboring PYRs from resting membrane potentials, that could be explained by differences in the amplitude of fast EPSCs evoked onto these cells (Figure 3F). Interestingly, as was observed at the network level, application of Atro. did not block the ripple activity in PYR current clamp recordings of single BLA PYRs, but instead increased its duration (Figure 3G). All ripple activity in the PYR membrane potential was completely blocked by Mec.

As the ACh-SWR was driven by nicotinic receptors, we hypothesized that the corresponding synaptic activity observed in PYR recordings would be driven by the same. To explore this, we examined the cholinergic response to single pulse light stimulation in BLA PYRs. In addition to the fast onset EPSCs and IPSCs (Figure 4A, inset), optogenetic ACh stimulation produced a direct slow outward current, as has been shown previously (36, 37). Not surprisingly, the fast onset event was blocked by mecamylamine (10 uM) while the late outward current was blocked by atropine (5 uM) (Figure 4B; Early area: Baseline = 1.31 ± 0.37; mec. = 0.15 ± 0.02; atro. = 0.06 ± 0.03, n = 11, N = 7; one-way ANOVA repeated measures, p = 4.74 x 10^-4^, F(_2,9_) = 20.14; Late area: Baseline = 10.66 ± 2.49; mec. = 9.74 ± 1.89; atro. = 0.14 ± 0.06, n = 11, N = 7; one-way ANOVA repeated measures, p = 4.46 x 10^-4^, F(_2,9_) = 11.29). The question then remained as to where these fast onset, highly summated EPSCs and IPSCs were originating. A primary option for EPSC generation is local PYR excitation, which could provide feedback excitation onto other BLA PYRs within the slice. This could be achieved by a direct nicotinic receptor-mediated excitation of BLA PYRs to spike threshold. However, application of CNQX blocked any significant effect by mecamylamine on the remaining PYR response (Figure 4C), indicating these cells do not exhibit a direct nicotinic current (Baseline area = 1.4 ± 0.34, CNQX = 0.24 ± 0.09, mec. = 0.21 = 0.08; n =5, N = 4; one way repeated measures ANOVA, p = 0.004, F(_1,4_) = 32.08). Alternatively, nicotinic receptors also function to increase glutamate release from pre-synaptic terminals from certain circuit pathways (31) and we have shown this to occur in BLA previously (32) on the millisecond timescale. This indicates that the increase in EPSCs in the BLA network was not due to direct excitation of BLA PYRs but instead likely due release of glutamate from cortical, among other, input pathways that express pre-synaptic nicotinic receptors at their axon terminals (32). Further, because these cell bodies from these inputs do not reside in the brain slice, these results suggest these EPSCs arise in an action potential-independent manner, whereby ACh binding on nicotinic receptors on these terminals results in vesicle release.

**Figure 4.**
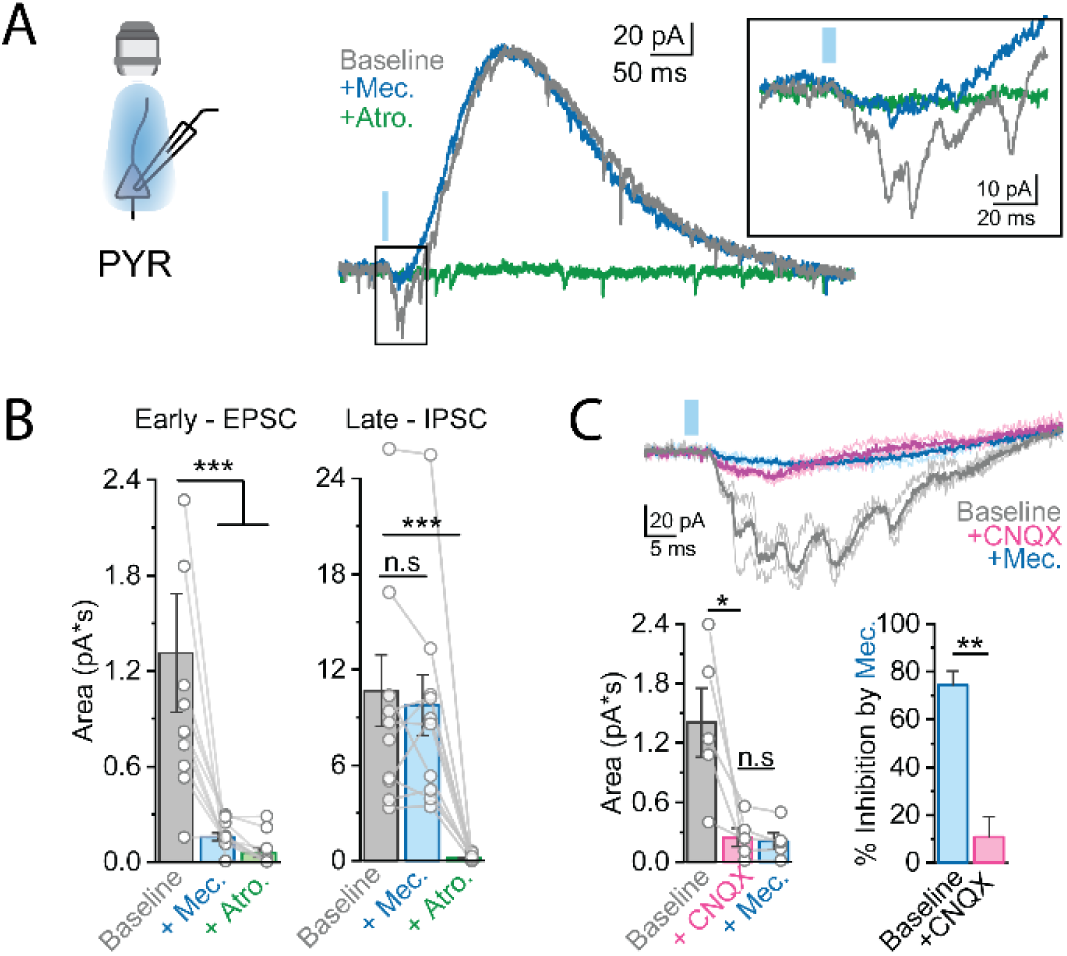
BLA PYRs do not show direct nicotinic-mediated currents and instead nicotinic-induced EPSCs arise from extra-amygdalar inputs. **A**. Voltage clamp recording of BLA PYRs during the entire cholinergic response following single pulse ACh stimulation shows a slow atropine-sensitive outward current that occurs after the fast barrage of events (inset, right) that are blocked by mecamylamine. **B**. Quantification of total area of the early events that are blocked by Mec. and the late event that is blocked by Atro. **C.** Application of CNQX abolishes the fast inward currents and subsequent application of Mec. no longer has any effect (Inhibition by Mec. in baseline = 71.93 ± 0.09%; inhibition by Mec. after CNQX = 10.80 ± 12.28%; two sample t-test, p = 0.002).

### Differential recruitment of BLA INs by acetylcholine nicotinic receptors

To this point we have established that nicotinic-receptor mediated SWRs in the BLA are generated in part through facilitation of glutamate release from external inputs to create a barrage of EPSCs. Next, we sought to explore the mechanism for the generation of the SWR-associated IPSCs. To do this, we performed recordings from putative BLA interneurons (INs), as could be distinguished by a series of electrophysiological parameters (high input resistance, low capacitance, characteristic spiking waveforms), in ChAT-ChR2 mice. In total, 36 putative IN recordings were used for this study. In response to the single pulse ACh stimulation, IN responses could be divided into three separate types based on their immediate (within 100 ms) response: no response (36%; 13/36 cells), jagged response (39%; 14/36 cells), or smooth response (25%; 9/36 cells) (Figure 5A). Both the jagged and smooth response types produced inward currents that in some cells could induce action potential firing from a resting membrane potential (Figure 5A, inset), indicating that single pulse ACh stimulation can induce local BLA IN firing, and thus induce IPSCs in local PYRs. Jagged and smooth responses differed in their rising kinetics. Jagged responses consisted of a summated inward current with many events contributing to a summed peak (Figure 5B), consistent with what looked like a barrage of EPSCs. Contrastingly, smooth responses exhibited a more linear onset to a single peak. These response types displayed distinct differences in their timing onset and peaks. Jagged responses exhibited a delayed onset from the light compared to the immediate smooth response (jagged = 11.26 ± 1.19 ms delay; smooth = 4.01 ± 0.35 ms; two-sample t-test, p = 9.57 x 10^-5^). Additionally, jagged responses took a significantly longer time to reach peak than the smooth responses (jagged = 28.18 ± 3.25 ms; smooth = 10.54 ± 2.39 ms; two-sample t-test, p = 8.21 x 10^-4^). These characteristic differences in these response types could be explained by differential sensitivity to blockade of AMPA receptors by CNQX (20 uM) (Figure 5C). Application of CNQX completely blocked the event in the jagged response group, but in cells with a smooth response the inward current was unaffected. The smooth response was also not blocked by NMDA or GABA receptor antagonists (APV, 50 uM; CGP, 1 uM; picrotoxin, 40 uM), indicating that this response is a direct post-synaptic nicotinic current in these cells. On the other hand, the jagged response reflects a barrage of nicotinic-mediated indirect AMPA EPSCs. These results show that single pulse cholinergic stimulation recruits BLA INs via two different nicotinic-mediated mechanisms that can be temporally separated; one through indirect glutamatergic events and one through a direct nicotinic current.

**Figure 5.**
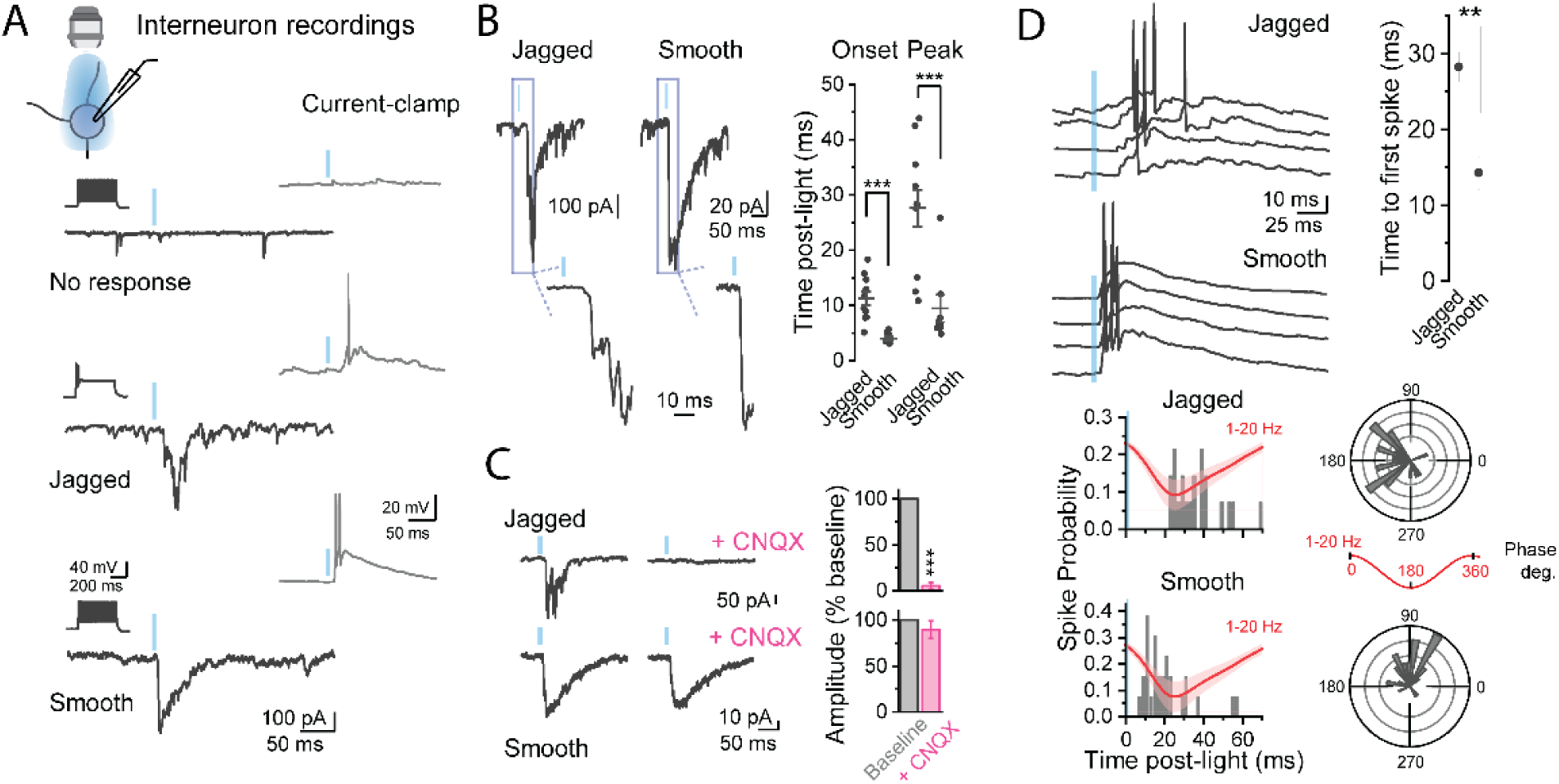
BLA interneurons are differentially recruited by acetylcholine nicotinic receptors to fire action potentials with different temporal profiles. **A.** Recordings from putative BLA interneurons (n = 36, N = 22) in response to single pulse ACh stimulation show three different response profiles: no response (36%), jagged response (39%), or smooth response (25%) as determined by voltage clamp and current clamp recordings. In a subset of these cells, stimulation could induce action potential firing (inset). **B.** Jagged and smooth responding cells could be distinguished based off the kinetics of the rise of their inward currents in response to ACh stimulation, where smooth responding cells showed an earlier current onset and time to peak response. **C.** The jagged and smooth responding cells could also be distinguished based off their sensitivity to CNQX, whereby CNQX blocked the response in the jagged cells but had no effect on smooth responding cells. **D.** Jagged and smooth responding cells also showed differences in the timing of their action potentials following ACh stimulation. Smooth responding cells showed a faster time to first action potential than jagged cells. Comparing the distribution of spike times across all trials shows (bottom) in comparison to the average sharp wave LFP (red trace) shows that while jagged cells show firing around the peak of the sharp wave, smooth cells tend to fire on the rising phase.

Because of this difference in response kinetics of BLA INs, we hypothesized that these cells would be recruited to spike at different times following ACh stimulation. To explore this, we performed current clamp recordings from the same INs at their resting membrane potential and recorded the timing of action potential firing. In total, 4 jagged response INs and 5 smooth response INs were induced to spike in response to ACh stimulation. Comparing the time to the first spike across the cells of each group shows that the jagged response group fired later after light stimulation than the early firing smooth responding cells (jagged = 28.25 ± 1.93 ms; smooth = 14.26 ± 2.13 ms; two sample t-test, p = 0.002). A comparison of the time course of spiking revealed a clear temporal difference in the probability of spiking in these different groups (Figure 4D) (jagged = 12 trials over 4 cells; smooth = 13 trials over 5 cells) when overlayed on the average SW of the ACh-SWR (red traces). Looking at the phase of action potential firing in these cells also shows a different preference on the average ACh-SWR (Figure 5E). These data indicate that these cells contribute differently to ACh-SWRs. Jagged response INs fired action potentials in close relation to the timing of LFP ripple peak (average ripple peak = 24.0 ± 1.2 ms) while smooth response INs were more closely aligned to ripple onset (average ripple onset = 14.8 ± 1.2 ms).

### Differential nicotinic receptor modulation of PV, CCK and SOM INs in the BLA

We next wanted to explore if BLA INs with these different ACh response types could be grouped into specific IN classes. Three large, mostly non-overlapping IN populations in the BLA are parvalbumin (PV), cholecystokinin (CCK), and somatostatin (SOM) INs. To explore the nicotinic receptor-mediated response in the different classes of BLA INs, targeted whole cell patch clamp recordings in fluorescently tagged cells from different transgenic mice lines were performed (PV-tdTomato, SOM-tdTomato, or CCK-Cre plus AAV-DIO-Dlx-mCherry virus injection, see Methods). We have previously shown these transgenic strategies to target distinct populations of BLA INs (35). In absence of ChR2 expression in ChAT neurons to optogenetically release ACh in these slices, brief (500 ms) focal puff application of ACh (1 mM) was applied within 100 microns of the recorded cell. Recordings from these different IN types revealed differences in fast (sub second onset) cholinergic responses (Figure 6A). The vast majority of PV INs did not exhibit direct fast responses to puff ACh (only 2/31 recorded cells, 6%), consistent with the lack of nicotinic response in these cells observed in other brain areas (38, 39). However, the majority of CCK interneurons (11/14 cells, 78%) responded with large direct nicotinic depolarization that could be blocked by mecamylamine (10 uM). Additionally, a subset of SOM INs also exhibited a large depolarization (5/21, 24%), which was also blocked by mecamylamine. These large nicotinic responses in CCK and SOM INs were not affected by blocking glutamate and GABA transmission, indicating they were a result of direct depolarizing nicotinic receptor response in these cells. These results indicate that a large population of CCK INs and a smaller population of SOM INs in the BLA exhibit nicotinic currents and likely contribute to the group of INs exhibiting smooth EPSCs in response to optogenetic cholinergic stimulation.

**Figure 6.**
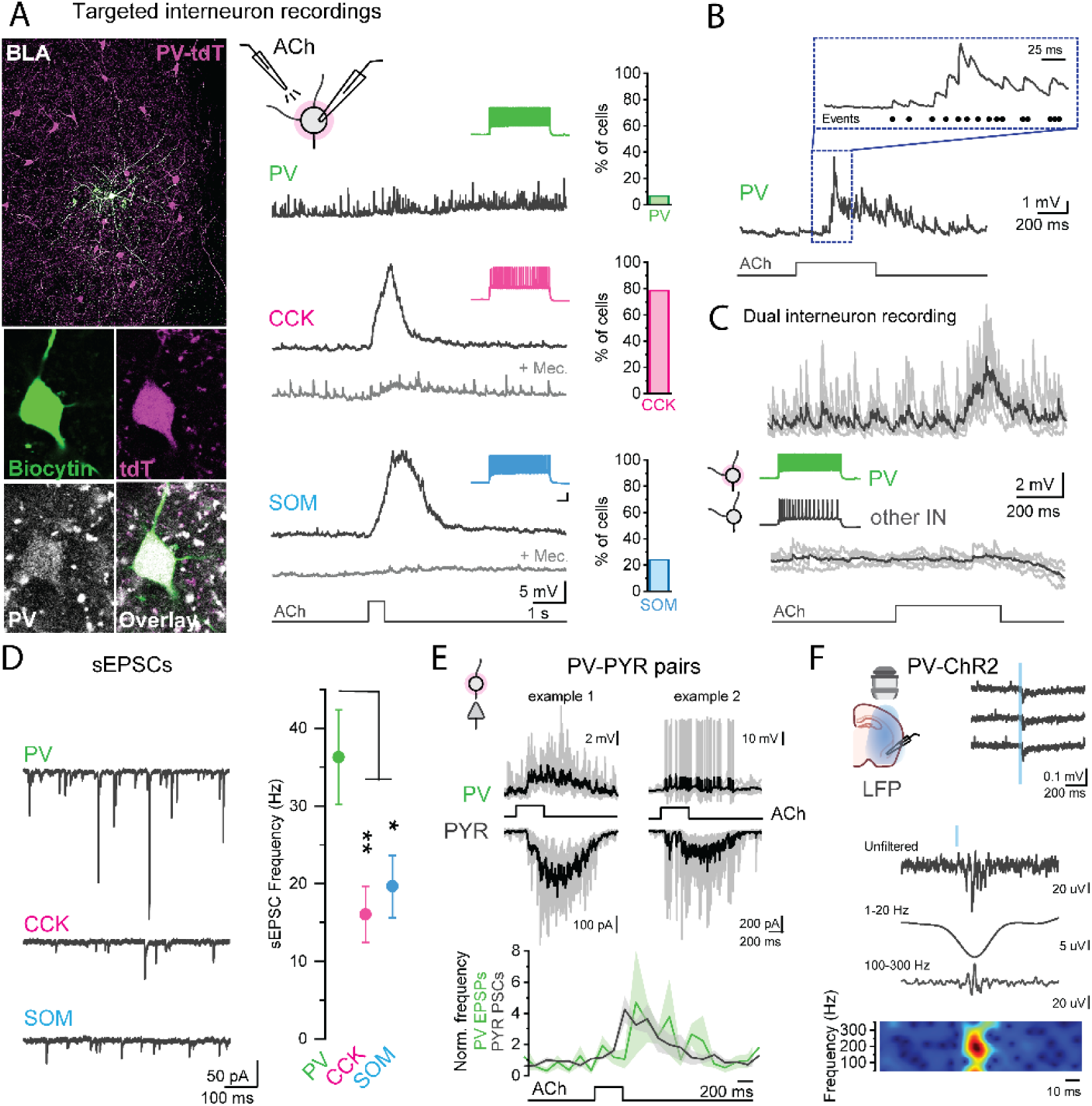
PV interneurons show increased EPSCs following focal acetylcholine application and can directly drive SWRs in the BLA. **A.** An example image of a targeted recording, as filled with biocytin in the recording electrode, of a PV IN in a PV-tdT slice. Focal puff application of ACh (1 mM) onto PV, CCK, and SOM interneurons elicits a fast response that can be blocked by mecamylamine in a subset of these cells. **B.** While most PV INs did not show a direct mecamylamine-sensitive depolarization, an example trace shows that a barrage of fast onset EPSP events could be detected. **C.** Dual recording from a PV IN and another BLA IN shows that this barrage of EPSPs could be present in PV INs specifically. **D.** Example traces of spontaneous EPSCs in these INs show that PV INs exhibit the highest frequency of events in baseline conditions. **E.** Two example recordings of PV and PYR pairs. PV INs receive a barrage of fast onset EPSPs, which sometimes induce PV spiking (right). The barrage of fast events in PV INs align in time with a barrage of post synaptic currents in PYRs (gray traces: single trials, black trace: mean response). **F.** Driving PV IN firing in slice preparations where PV INs express ChR2 shows that single pulse stimulation of these cells can induce network events in the LFP recording (top, example traces). A closer examination of these events shows they closely resemble sharp-wave ripple activity, indicating that PV INs can drive SWRs in BLA slices.

**Table 1.** Statistical table.

| Figure | Parameter | Sample Size (n, N) | Primary Statistical Test | Post-hoc Test | Comparison | P | F/t statistic |
| --- | --- | --- | --- | --- | --- | --- | --- |
| 1B | LFP: sharp wave and ripple time course - Onset | 22, 15 (sharp wave)<br>12, 12 (ripple) | Paired t-test | | Sharp wave vs ripple | $1.73 \times 10^{-4} ***$ | $t(11) = -5.54$ |
| 1B | LFP: sharp wave and ripple time course - Peak | 22, 15 (sharp wave)<br>12, 12 (ripple) | Paired t-test | | Sharp wave vs ripple | 0.84 | $t(11) = -0.20$ |
| text | ACh-SWR in male and female mice | 14, 10 (male)<br>8, 5 (female) | Two sample t-test | | SW amplitude: male vs female | 0.91 | $t(20) = -0.11$ |
| | | | | | SW duration: male vs female | 0.97 | $t(20) = 0.04$ |
| | | | | | Ripple duration: male vs female | 0.27 | $t(10) = -1.17$ |
| 1E | Low vs high frequency cholinergic stimulation | 8, 5 | One way repeated measures ANOVA | | Low frequency all pulses: SW amplitude | $7.78 \times 10^{-11} ***$ | $F(4,11) = 263.00$ |
| | | 5, 4 | | | Low frequency all pulses: ripple amplitude | $4.058 \times 10^{-7} ***$ | $F(4,4) = 2717.51$ |
| | | 15, 9 | | | High frequency all pulses: SW amplitude | 0.144 | $F(4,4) = 3.17$ |
| | | 8, 6 | | | High frequency all pulses: ripple amplitude | 0.079 | $F(1,4) = 89.82$ |
| 2A | Sharp wave amplitude: atro | 8, 6 | Paired t-test | | Baseline vs atro | 0.24 | $t(7) = 1.26$ |
| | Sharp wave amplitude: mec | 11, 7 | Paired t-test | | Baseline vs mec | $1.31 \times 10^{-4} ***$ | $t(10) = 26.15$ |
| | Sharp wave amplitude: CNQX | 7, 4 | Paired t-test | | Baseline vs CNQX | 0.024* | $t(6) = 3.01$ |
|  | Sharp wave amplitude: Picrotoxin | 6, 4 | Paired t-test |  | Baseline vs picrotoxin | 0.028* | t(5) = 3.06 |
| 2B | Ripple duration: atro | 3, 3 | Paired t-test |  | Baseline vs atro | 0.13 | t(2) = -2.48 |
|  | Ripple duration: mec | 3, 3 | Paired t-test |  | Baseline vs mec | 0.002** | t(2) = 21.40 |
| 2E | Multiunit frequency: 0-50 ms |  | Paired t-test |  | Baseline vs atro | 0.355 | t(3) = -1.089 |
|  |  |  | Paired t-test |  | Baseline vs mec | 0.031* | t(4) = 3.253 |
|  |  |  | Paired t-test |  | Baseline vs picro | 0.026* | t(5) = 3.116 |
|  |  |  | Paired t-test |  | Baseline vs CNQX | 0.010** | t(5) = 3.925 |
| 2E | Multiunit frequency: 50-100 ms |  | Paired t-test |  | Baseline vs atro | 0.019* | t(3) = -4.550 |
|  |  |  | Paired t-test |  | Baseline vs mec | 0.044* | t(4) = 2.282 |
|  |  |  | Paired t-test |  | Baseline vs picro | 0.011* | t(5) = 3.952 |
|  |  |  | Paired t-test |  | Baseline vs CNQX | 0.048* | t(5) = 2.318 |
| 2F | Effects of physostigmine on ACh-SWRs: SW amplitude | 22, 15 (no physo.)<br>27, 17 (with physo) | Two sample t-test |  | No physo. vs physo. | 0.627 | t(47) = -0.488 |
|  | SW duration | 22, 15 (no physo.)<br>27, 17 (with physo) | Two sample t-test |  | No physo. vs physo. | 0.452 | t(47) = -0.758 |
|  | Ripple duration | 12, 8 (no physo.)<br>8, 6 (with physo) | Two sample t-test |  | No physo. vs physo. | 0.388 | t(18) = -0.883 |
|  | Multiunit | 20, 14 (no physo.)<br>24, 15 (with physo) | Two sample t-test |  | No physo. vs physo | 0.909 | t(42) = 0.114 |
| 3A | PYR voltage clamp: EPSCs vs IPSCs onset | 22, 15 | Two sample t-test |  | EPSCs vs IPSCs | 1.97 x 10<sup>-5***</sup> | t(47) = -4.74 |
| 3A | PYR voltage clamp: EPSCs vs IPSCs peak | 22, 15 | Two sample t-test |  | EPSCs vs IPSCs | 0.07 | t(47) = 1.82 |
| 3G | PYR current clamp: ripple duration | 7, 3 | One way repeated measures ANOVA |  | Overall | 4.69 x 10<sup>-5***</sup> | F(2, 5) = 132.13 |
| | | | | Tukey | Baseline vs Atro, | $2.05 \times 10^{-7***}$ | |
| | | | | Tukey | Baseline vs Mec. | $5.06 \times 10^{-5***}$ | |
| 4B | PYR voltage clamp cholinergic response: Early | 11, 7 | One way repeated measures ANOVA | | Overall | $4.74 \times 10^{-4***}$ | $F(2, 9) = 20.14$ |
|  |  |  |  | Tukey | Baseline vs Mec | 0.003** |  |
|  |  |  |  | Tukey | Mec vs Atro | 0.94 |  |
| 4B | PYR voltage clamp cholinergic response: Late | 11, 7 | One way repeated measures ANOVA | | Overall | $4.46 \times 10^{-4***}$ | $F(2,9) = 11.29$ |
|  |  |  |  | Tukey | Baseline vs Mec | 0.87 |  |
| | | | | Tukey | Mec vs Atro | $3.57 \times 10^{-5***}$ | |
| 4C | PYR voltage clamp cholinergic response: Early EPSCs | 5, 4 | One way repeated measures ANOVA | | Overall | 0.004** | $F(1,4) = 32.08$ |
|  |  |  |  | Tukey | Baseline vs CNQX | 0.015* |  |
|  |  |  |  | Tukey | CNQX vs Mec | 0.992 |  |
| 4C | Inhibition by mec | 11, 7 (baseline)<br>5, 4 (+CNQX) | Two sample t-test | | Baseline vs +CNQX | 0.002** | $t(13) = -3.89$ |
| 5B | Jagged and Smooth responding interneurons: Response onset | 11, 8 (jagged)<br>8, 7 (smooth) | Two sample t-test | | Jagged vs smooth | $9.57 \times 10^{-5***}$ | $t(17) = 5.06$ |
| 5B | Jagged and Smooth responding interneurons: Response peak | 11, 8 (jagged)<br>8, 7 (smooth) | Two sample t-test | | Jagged vs smooth | $8.21 \times 10^{-4***}$ | $t(17) = 4.05$ |
| 5D | Jagged and Smooth responding interneurons: Time to first action potential | 4, 4 (jagged)<br>5, 4 (smooth) | Two sample t-test | | Jagged vs smooth | 0.002 | $t(7) = 5.36$ |
| 6D | Spontaneous<br>EPCS<br>frequency | 11, 7 (PV)<br>16, 7 (SOM)<br>16, 3 (CCK) | One way<br>ANOVA | | Overall | 0.009 | $F(2,40) = 5.21$ |
|  |  |  |  | Bonferroni | PV vs SOM | 0.044 |  |
|  |  |  |  | Bonferroni | PV vs CCK | 0.010 |  |
|  |  |  |  | Bonferroni | SOM vs CCK | 1 |  |

While the majority of PV INs did not show direct nicotinic currents, in many cells we observed a quick onset barrage of EPSPs (Figure 6B), closely matching the jagged response phenotype from optogenetically released ACh. Even in paired recordings of a PV IN and a neighboring non-PV IN, application of ACh could induce an EPSP barrage in the PV IN (Figure 6C) and not the neighboring IN, suggesting that these cells are preferentially exhibiting this response. In the BLA, as in other cortical structures, PV INs receive an especially dense innervation from both local and long range projections (40, 41), that exceeds that of other IN types. In agreement, recording spontaneous EPSCs (sEPSCs) from PV, CCK, and SOM INs in our preparations revealed that PV INs received the highest frequency of sEPSCs at a rate that almost doubled the other IN types (PV sEPCS frequency = 36.28 ± 6.09 Hz; CCK = 16.03 ± 3.64 Hz; SOM = 19.64 ± 4.00 Hz; one way ANOVA, p = 0.009) (Figure 6D). This positions these cells to play an important role in glutamatergic-mediated network events. Paired recordings of PV and PYRs show that PV EPSPs aligned in time with the increase in PYR PSCs (Figure 6E). Indeed, in slice preparations of PV-Cre mice where PV INs virally express ChR2, brief (1 ms) light stimulation of PV INs induced a SWR event of the LFP (Figure 5F). These SWRs had immediate onset (1.41 ± 0.08 ms after beginning of light stimulation; 3 slices, 2 animals) and exhibited similar characteristics to ACh-SWRs (duration = 185.83 ± 9.01 ms). This indicates that PV IN activity can directly induce BLA SWRs, in agreement with our prior findings (35). Together, these results show that PV INs are critical drivers of ACh-SWRs.

## Discussion

The results of the present study outline a previously unknown mechanism by which low frequency cholinergic stimulation can induce SWRs in BLA through engagement of nicotinic receptors. These ACh-SWRs closely mimic the time course, frequency, and underlying local network behavior of those observed *in vivo* and *in vitro* in both the BLA and hippocampus. Muscarinic receptors were not involved in the induction of ACh-SWRs, but instead constrained their duration to a tight temporal window following cholinergic stimulation. At the synaptic level, nicotinic receptors induce ACh-SWRs through recruitment of ripple-frequency glutamatergic and GABAergic events. Excitation is induced through a pre-synaptic facilitation of glutamate release from extra-amygdala inputs that excites both BLA PYRs and distinct subtypes of INs, leading to ripple-associated inhibition. Subclasses of both CCK and SOM INs are directly activated by nicotinic receptors and can fire fast-onset action potentials. On the other hand, PV INs, which receive the most glutamatergic inputs of BLA INs, rarely showed a direct nicotinic current but instead uniquely displayed a barrage of EPSCs that was associated with PYR ripple-associated activity. In line with this, direct PV IN activation is sufficient for SWR generation in the BLA in baseline states. In total, this study establishes a novel mechanism for SWR generation in the BLA in which the cholinergic system extends beyond a purely neuromodulatory role to directly induce SWRs.

### Nicotinic receptor induction of ACh-SWRs

These data shed new light on how basal forebrain cholinergic neurons and cholinergic tone modulate SWRs. In the hippocampus, ACh can suppress SWRs through a muscarinic receptor mechanism, likely by inhibiting glutamate release between PYRs (25–27, 42). These studies used broad application of ACh agonists or large optogenetic cholinergic stimulations, however, and did not explore effects of nicotinic receptors or sparse cholinergic stimulation. Nicotinic receptors are known to facilitate glutamate release from presynaptic terminals (31) on millisecond timescales (32, 43), and thus are well positioned to induce SWRs, which are reliant on fast glutamatergic transmission. However, nicotinic receptors are uniquely vulnerable to receptor desensitization and thus exploration of their effects can be difficult to interpret with broad agonists (44). To this end, optogenetic tools allow for synaptic release of ACh at physiological concentrations and offer better insight into *in vivo* ACh transmission. Likewise, sparse or low frequency cholinergic stimulation, as used in the present study, is similar to cholinergic activity observed during SWS or periods of quiet wakefulness when SWRs occur most frequently (19). The ability of sparse or low frequency cholinergic activity to induce SWRs in the BLA likely reflects the stronger cholinergic innervation of BLA than hippocampus. Indeed, we have previously shown that in BLA ACh release can potentiate glutamate release on the timescale of an individual synaptic event (32). These findings combined with the current study further suggest that cholinergic signaling can serve precise, computational roles in the BLA network.

Synaptic acetylcholine can bind both fast ionotropic nicotinic receptors (millisecond onset) and slow metabotropic muscarinic receptors (100s of millisecond onset). Intriguingly, this allows for a short window by which single event ACh release can bind nicotinic receptors to modulate network activity unopposed by competing muscarinic influence. Indeed, this is consistent with our results in the present study. We have previously shown that prolonged high frequency cholinergic stimulation reduces low frequency network activity via a muscarinic receptor mechanism (35), indicating that as in the hippocampus, muscarinic receptors can inhibit BLA SWRs. In the present study, single pulse stimulation of cholinergic fibers induced an immediate nicotinic receptor-mediated ACh-SWR. This event was prolonged by inhibiting muscarinic receptors, suggesting that even at that short time scale muscarinic receptors can act to inhibit SWR activity, and leaves a short window whereby nicotinic receptors can dominate in their circuit influence. Further, reliable ACh-SWR induction only occurred during slow frequency periodic basal forebrain cholinergic neuron firing, as is observed during immobility and SWS, as opposed to 5 Hz stimulation, as is seen during active exploration or REM sleep. This positions basal forebrain neurons to switch between network states in the BLA permissible to SWR generation and offers a mechanism by which the basal forebrain cholinergic neurons could drive emotional memory consolidation during SWS. Further, these results call for an update in the field’s view of neuromodulatory systems in SWRs. Depending on the receptor subtypes recruited or the mode of firing of these systems, it is possible they can cause temporally precise, direct induction of SWRs.

### Parallels between ACh-SWRs in the BLA and SWRs in the hippocampus

Interestingly, the interplay of nicotinic and muscarinic receptors on BLA SWRs closely mimics several aspects of hippocampal SWRs. In the hippocampus, SWRs in CA1 depend on an influx of highly synchronized excitatory input from CA3 (3). This initiates the SWR event which persists for 50-150 ms and is then terminated by a large hyperpolarizing reflection that is likely a K+ mechanism (45), notably a GIRK, that is believed to be produced from an intrinsic circuit event. In parallel, in the BLA nicotinic receptors induce synchronized excitation through glutamate release from extra-amygdalar axon terminals. The single ACh stimulus can act on multiple axon terminals to produce a highly synchronized ripple-frequency barrage of EPSCs across populations of BLA cells, analogous to the synchronized CA3 discharge to CA1 (1). The simultaneous action of ACh on these pre-synaptic receptors is an action potential-independent mechanism for these pre-synaptic neurons and bypasses the need for a synchronized discharge in upstream regions. Subsequently, muscarinic receptors induce a large GIRK-mediated hyperpolarization of BLA PYRs that peaks between 100-200 ms after the ACh stimulus, providing an analogous event to terminate the SWR as is observed in the hippocampus. Indeed, our results show that within this 100 ms time window following ACh stimulation, blocking muscarinic receptors can prolong the increase in multiunit activity and ripple-frequency membrane potential oscillation during this period. This is an interesting mechanism by which ACh signaling acting through its two different types of receptors can work in a competing mechanism to induce a temporally specific network ACh-SWR in the BLA that is similar to the traditional mechanism of hippocampal SWRs. Further, this establishes that nicotinic and muscarinic receptors exert different effects on SWRs and allows for the mode of firing in basal forebrain cholinergic neurons to control these SWRs in a frequency-dependent manner.

### Differential recruitment of BLA interneurons to the ACh-SWR

The ACh-SWRs in the BLA recruited distinct populations of local INs through a direct and indirect mechanism, resulting in differential timing of action potential firing across INs. This suggests differential roles for specific interneurons during BLA SWRs, as has been noted in hippocampus (20, 46). ACh puff application suggests that a large population of CCK and a smaller population of SOM INs are directly depolarized by nicotinic receptors and could be induced to fire quick onset action potentials. This is the first insight into the firing of these BLA IN populations during SWRs. In the hippocampus, it is still not entirely clear the role that these IN populations play in SWRs, but they tend to show only little action potential firing (46–48). Similar to other areas, BLA PV INs receive large amounts of excitatory innervation (49) and could receive a barrage of summating EPSCs induced indirectly by nicotinic receptors. These cells showed firing at times where peak ripple activity was observed, consistent with activity in these cells in hippocampal and BLA SWRs (18, 46). This suggests that the SWRs induced in the BLA show similar cell recruitment to those in the hippocampus. Consistent with this, optogenetic activation of PV INs in the BLA was sufficient for SWR induction, as observed in the hippocampus (50). Future work will be needed to resolve the precise function of different IN classes in ACh-SWRs, specifically, looking at how the temporal evolution of IN recruitment by ACh shapes this event.

### Potential role of ACh-SWRs in emotional memory consolidation

The results of the present study have intriguing implications for BLA memory consolidation, which is known to occur in the BLA to induce plastic changes in the circuit (12, 51–53). Despite this, there is not a great understanding for the mechanism of memory consolidation in the BLA (54). In the hippocampus, SWRs are associated with firing of PYRs previously active during a past experience (8, 55). Indeed, inhibiting these specific PYRs after learning can block memory consolidation (56). The reactivation of specific PYR sequences is reliant on NMDA receptors during learning, but not after (10), indicating that active cells are likely “tagged” during encoding to be selectively reactivated offline during consolidation. Indeed, the population of PYRs that are activated during SWRs have an unbalanced excitation/inhibition ratio favoring excitation (57), indicating that these PYRs show short term potentiation to their excitatory inputs and/or have undergone plastic changes within local IN circuitry, contributing to their reactivation during offline SWRs. In hippocampal brain slices, long-term potentiation (LTP) protocols can induce increased SWRs (21) in the network, further indicating a role of excitatory potentiation in altering SWR activity.

Our results show within a given preparation that some PYRs are depolarized while others are hyperpolarized during ACh-SWRs. We hypothesize that this is likely driven by the balance of EPSCs/IPSCs induced by ACh in these cells. Indeed, we noticed a large spread in the amplitude of these events across recordings PYRs. This selective firing is consistent with what occurs *in vivo*, which could be explained by differences in synaptic weights in PYRs from the underlying circuitry (58). Along these lines, fear learning results in potentiated cortical inputs to BLA PYRs (59) as well as reduced PV-mediated inhibition (60). Increased glutamatergic drive and decreased PV mediated inhibition onto these cells would enhance the excitation-inhibition balance and may offer a “tagging” mechanism for selective reactivation of these PYRs during ACh-SWRs, especially as cortical inputs express nicotinic receptors on their axon terminals. On the other hand, following fear extinction, previously fear-activated cells now receive an increase in PV+ terminals around their cell body (61). This would result in a decrease in the excitation/inhibition ratio onto these PYRs and now might inhibit these cells during subsequent ACh-SWRs at the expense of newly activated fear extinction neurons. In our hands, ACh-SWRs showed a remarkable consistency across repeated stimulations, lending to the hypothesis that these events could be replaying precise ensemble activity. Future studies using brain slices of mice that have recently undergone fear learning are needed to explore this intriguing idea.

In total, these hypotheses position nicotinic receptors as important drivers of BLA memory consolidation. Nicotinic receptor agonists have been implicated in promoting memory consolidation when administered after learning (62), but show differences across brain areas (63, 64). There have not been any studies exploring the direct effects of nicotinic receptors in the BLA *after* learning and their role in memory consolidation in BLA circuits, but the results from the present study outline a potentially important role for these receptors in emotional memory consolidation via induction of BLA SWRs.

## Materials and Methods

### Animals

All animal procedures used in this study were first approved by the University of South Carolina Institutional Animal Care and Use Committee under guidelines outlined in the Health Guide for the Care and Use of Laboratory Animals from the National Institution of Health. All mice were purchased from The Jackson Laboratory and maintained at the University of South Carolina’s lab animal resource facility on a 12-hour light/dark cycle, where they were provided *ad libitum* food and water. Mice were housed with littermates at appropriate density for all studies. All mice used in this study were between 5 and 16 weeks old and included both male and female mice. Differences in male and female data were explored for select data sets and reported. For selective optogenetic activation of cholinergic neurons, our lab used the following breeding pairs to create the a double transgenic line which we termed ChAT-ChR2: female Chat-Cre mice (B6;129S6-Chat ^tm2(cre)Lowl^/J, stock number #006410, The Jackson Laboratory) crossed with male Ai32 mice (B6; 129S-Gt(ROSA)^26SOR tm21(CAG-COP4*H134R/EYFP)Hze^ /J, #012569, The Jackson Laboratory). These mice have been used in previous studies in our lab (32, 35) and have been shown to express ChR2 specifically in ChAT neurons. For targeted IN patch clamp recordings, our lab developed multiple double transgenic lines: PV-tdTomato mice - created by crossing female PV-Cre (B6;129P2-*Pvalb^tm1(cre)Arbr^*/J,#008069, The Jackson Laboratory) with male Ai14 (B6.Cg-*Gt(ROSA)26Sor^tm14(CAG-tdTomato)Hze^*/J,#007914, The Jackson Laboratory) and SOM-tdTomato mice - created by crossing female SOM-Cre (*Sst^tm2.1(cre)Zjh^*/J,#013044, The Jackson Laboratory) with male Ai14. For CCK IN recordings, due to the non-specific expression of CCK in both PYRs and INs, CCK Ins were instead targeted with an interneuron specific viral promoter injected into the BLA in the CCK-Cre mouse line (C57BL/6-Tg(Cck-cre)CKres/J,#011086, The Jackson Laboratory) (65).

### Stereotaxic viral surgeries

Stereotaxic viral injections were performed in mice that were 6-12 weeks of age. Mice were first anesthetized with isoflurane and then secured in the stereotaxic configuration using ear cups (Stoelting). For all experiments, viral injections were targeted to the BLA of both hemispheres using the following coordinates from bregma: anterior/posterior -1.4 mm, medial/lateral ±3.2 mm, dorsal/ventral -5.2mm. For channelrhodopsin experiments in PV interneurons, PV-Cre mice were injected with 200 nl of AAV5-EF1a-DIO-hChR2(H134R)-eYFP (UNC Vector Core) in each hemisphere. For experiments targeting CCK interneuron recordings, CCK-Cre mice were injected with 350 nl of AAV5-hDlx-Flex-dTomato-Fishell_7 (Addgene) in each hemisphere. Surgical incisions were closed with GLUture adhesive (Abbott Laboratories) following surgery and monitored during recovery. Mice were allowed at least 3 weeks before use in brain slice electrophysiological experiments to enable sufficient viral expression in the cells.

### Brain Slice Preparation

For brain extraction, mice were first placed under deep isoflurane anesthesia. Brains were quickly removed and fully submerged directly into a small container with ice-cold cutting solution containing (in mM): 110 choline chloride, 2.5 KCl, 25 NaHCO_3_, 1.25 NaH_2_PO_4_, 20 glucose, 5 MgSO_4_, and 0.5 CaCl_2_ and vigorously bubbled with 95% O_2_ and 5% CO_2_. Brains were kept in this solution long enough for the whole brain to cool, or approximately 1 minute. From here, the brains were transferred to a Leica VT1000S vibratome, the cerebellum blocked, and the remaining brain glued securely onto a cutting platform. For LFP recordings, coronal slices were cut at 400-500 microns to preserve maximal circuitry. For whole cell patch clamp recordings, coronal slices of 300 microns were made. Slices were placed in a fresh solution after cutting containing (in mM): 125 NaCl, 2.7 KCl, 25 NaHCO_3_, 1.25 NaH_2_PO_4_, 10 glucose, 5 MgSO_4_, and 0.5 CaCl_2_, superfused with 95% O_2_ and 5% CO_2_ at 34°C (pH 7.3; 300-310 mOsm). Upon the slices being placed in the incubation chamber, this incubation artificial cerebral spinal fluid (ACSF) was held at 34°C for 30-45 minutes before being allowed to gradually cool to room temperature. Slices were allowed at least 45 minutes to recover before being used for electrophysiology recordings.

### Brain Slice Electrophysiology Recordings

#### LFP Recordings

Slices 400-500 µm thick were transferred to a recording chamber containing an ACSF solution containing (in mM): 125 NaCl, 3.3 KCl, 1.25 NaH_2_PO_4_, 25 NaHCO_3_, 10 glucose, 2 CaCl_2_ and 1 MgSO_4_ (pH 7.3; 300-310 mOsm) for experimental procedures. The recording solution was perfused over the slice at an accelerated rate of 7-12 mL/minute to achieve maximal slice activity and health and was continuously bubbled with 95% O2/5% CO2 gas. All recordings were done at 32 - 34°C. Borosilicate glass electrodes with a resistance of 1-2 MOhms were used for recordings. Electrodes were placed into the BLA near a collection of cell bodies, as visualized with infrared-differential interference contrast optics (Olympus BX51WI) through a 40X water immersion lens. Recordings did not start until at least one minute after the electrode was placed to avoid confounding activity induced by the electrode disrupting the tissue. Optogenetic stimulation of cholinergic fibers was achieved by 1-2 millisecond duration pulses of 490 nm light delivered through the 40X objective lens directly over the recording electrode (pE-4000, CoolLED). Light stimulation was given once every 60-90 seconds, unless otherwise noted. Recordings were filtered at 2 kHz and amplified by 1000-2000X (Multiclamp 700A amplifier, Digidata 1440A A-D board, Molecular Devices). For SWR detection, traces were bandpass filtered at different frequencies: SW = 1-20 Hz, ripple = 100-300 Hz, multi-units = 500-5000 Hz. SWR analysis was performed on the different filtered signals for each trace. Ripple presence was assigned only if waveform amplitude significantly deviated from the baseline average. For visualization of waveform spectrograms, short-time Fourier transforms (STFTs) were created from 300-600 millisecond time bins around cholinergic light stimulation. For multi-units, pClamp threshold detection was utilized to detect unit activity and all units were manually inspected to ensure the waveform resembled unit activity and not electrical noise. Multi-unit frequency in response to light stimulation was binned into 10 ms time bins for analysis.

#### Whole Cell Recordings

Slices of 300 µm were used for whole cell experiments for visual clarity of single cells. Recordings were performed in the same chamber and set up as LFP recordings, but with a slower perfusate rate (2-4 mL/minute) and a recording ACSF solution with lower extracellular K^+^ (whole cell recording ACSF (in mM): 125 NaCl, 2.7 KCl, 1.25 NaH_2_PO_4_, 25 NaHCO_3_, 10 glucose, 2 CaCl_2_ and 1 MgSO_4_ - pH 7.3; 300-310 mOsm). BLA PYRs or INs were targeted for recordings based off their cell body size and confirmed by measuring electrophysiological parameters. All cells were recorded with borosilicate glass electrodes with an input resistance of 4-6 MOhms. Recording electrodes were filled with the following internal solutions. For PYR voltage clamp and IN voltage and current clamp recordings: (in mM) 135 K-gluconate, 5 KCl, 10 HEPES, 2 MgCl_2_, 2 MgATP, 0.3 NaGTP, and 0.5 EGTA (pH 7.3, 290 mOsm). For PYR neuron current clamp recordings: (in mM) 137 K-gluconate, 3 KCl, 10 HEPES, 2 MgCl_2_, 2 MgATP, 0.3 NaGTP, and 0.5 EGTA (pH 7.3, 290 mOsm).

#### Drug Application

For pharmacological experiments, drugs were applied to the recording ACSF solution and allowed at least two minutes to perfuse onto the slice before measurements were made. The following drugs were used: 6-cyano-7-nitroquinoxaline-2,3-dione disodium salt (CNQX, 20 µM, Hello Bio), picrotoxin (50 µM, Hello Bio), atropine (5 uM, Abcam, mecamylamine (20 uM, Millipore Sigma), and physostigmine (1 uM, Hello Bio). For recordings where acetylcholine was focally applied, a recording borosilicate glass electrode was filled with the recoding ACSF solution containing 1 mM acetylcholine chloride (Millipore Sigma) and applied within 100 microns of the recorded cell using a brief pulse (500 ms) from a picospritzer.

## Data Analysis

pClamp 10 software was used for initial processing and analysis of electrophysiology data. Further analysis and statistical tests were performed using OriginPro software. In all figures, data is presented as mean ± SEM unless otherwise noted, such as for box and whisker plots where the individual data points are plotted and lines at the 25^th^ and 75^th^ percentiles, median designated by the middle solid line, and the mean represented as a square data point. For determining significance in statistical tests, *α* < 0.05 was used and the following designations were assigned for the plots in the figures: * = *p* < 0.05, ** = *p* < 0.01, and *** = *p* < 0.001.

## Acknowledgments

We thank all members of the Mott laboratory for their comments on this project. This work was supported by National Institute of Mental Health R01MH104638 to D.D.M. and A.J.M. and R01MH131808 to D.D.M.; University of South Carolina VP for Research ASPIRE2 Award to D.D.M.; University of South Carolina VP for Research Support to Promote the Advancement of Research and Creativity (SPARC) Research Grant to J.X.B-P.

## Author Contributions

Designed research: J.X.B-P and D.D.M. Performed research: J.X.B-P, G.C.J and J.W.W. Analyzed data: J.X.B-P. Wrote the paper: J.X.B-P and D.D.M. Edited the paper: J.X.B-P, and D.D.M. Obtained funding for the research: J.X.B-P and D.D.M.

